# MoBiFC: development of a modular bimolecular fluorescence complementation toolkit for the analysis of chloroplast protein-protein interactions

**DOI:** 10.1101/2021.03.01.433373

**Authors:** Florent Velay, Melanie Soula, Marwa Mehrez, Stefano D’Alessandro, Christophe Laloi, Patrice Crete, Ben Field

## Abstract

The bimolecular fluorescence complementation (BiFC) assay has emerged as one of the most popular methods for analysing protein-protein interactions (PPIs) in plant biology. This includes its increasing use as a tool for dissecting the molecular mechanisms of chloroplast function. However, the construction of chloroplast fusion proteins for BiFC can be difficult, and the availability and selection of appropriate controls is not trivial. Furthermore, the challenges of performing BiFC in restricted cellular compartments has not been specifically addressed. Here we describe the development of a flexible modular cloning-based toolkit (MoBiFC) for chloroplast BiFC and proximity labelling using synthetic biology principles. The approach facilitates the cloning process for chloroplast-targeted proteins, allows robust ratiometric quantification, and the toolkit comes with model positive and negative controls. Our study also highlights many potential pitfalls including the choice of fluorescent protein (FP) split, negative controls, cell type, and reference FP. Finally, we provide an example of how users can enrich the toolset by providing functional proximity labelling modules, and we discuss how MoBiFC could be further improved and extended to other compartments of the plant cell.

## Introduction

The characterisation of protein protein interactions (PPIs) is important for understanding the assembly of macromolecular machines and deciphering the signal transduction pathways that are required for regulating the growth, development, and acclimation of plants in a constantly changing environment. A wide range of methods are used to study PPIs, including yeast two-hybrid, affinity purification, and protein-fragment complementation assays (Lampugnani et al., 2018; Struk et al., 2019). The bimolecular fluorescence complementation (BiFC) or split-fluorescent protein (FP) assay has emerged as the most popular protein-fragment complementation method in plant biology (Miller et al., 2015; Kudla and Bock, 2016; Struk et al., 2019). BiFC relies on assembly of a stable FP upon the interaction between two proteins of interest (POIs) each fused to separate non-fluorescent fragments of the FP (Romei and Boxer, 2019). The popularity and power of BiFC lies in its relative simplicity, in vivo nature, and ability to capture weak or transient PPIs. The first BiFC assay was performed using a GFP split in mammalian cells (Hu et al., 2002). Subsequently, many different splits and GFP variants were developed with the aim of improving signal strength and reducing non-specific complex assembly (Kudla and Bock, 2016; Struk et al., 2019). Further innovations followed, including the development of vector systems allowing the co-expression of the POIs and reference FPs from the same construction (Grefen and Blatt, 2012; Gookin and Assmann, 2014), the use of super-resolution compatible FPs, and multicolour or trimolecular complementation assays that allow the simultaneous visualization of multiple protein complexes (Struk et al., 2019). While BiFC is an attractive and powerful technique, the stable nature of the assembled FP means that particular attention must be paid to the selection of appropriate controls (Kodama and Hu, 2012; Gookin and Assmann, 2014; Horstman et al., 2014; Kudla and Bock, 2016). The gold-standard negative control is a BiFC assay using a mutated POI lacking its interaction domain.

In plants there are reports of successful BiFC assays in multiple cellular compartments, including the cytosol, nucleus (Bracha-Drori et al., 2004), chloroplast (Maple et al., 2005; Citovsky et al., 2006), and more recently in the mitochondria (Zhang et al., 2017; Shin et al., 2020). However, despite the growing number of studies reporting organellar PPIs by BiFC only a handful use gold-standard negative controls (Maple et al., 2007; Wang and Blumwald, 2014; Ramos-Vega et al., 2015; Hong et al., 2020; Ouyang et al., 2020; Sun et al., 2020) and many fall far short. For example, a surprising number of studies lack any negative control, or use the expression of a cytosolic FP fragment as a negative control for the interaction between chloroplastic POIs. In the latter case, the negative control will not reflect non-specific assembly because, without a chloroplast transit peptide, the free FP fragment cannot encounter the POI in the chloroplast. In one study, a free FP fragment was addressed to the chloroplast for the negative control, which, if correctly targeted, would provide a better indication of the level of non-specific assembly (Zhang et al., 2016). In addition, no study has specifically addressed whether the small volume of compartments in the chloroplasts and mitochondria can affect the outcomes of BiFC assays. Indeed, because BiFC fusion proteins are usually highly expressed, their concentration in restricted cellular compartments will increase the frequency of random protein-protein collisions and could lead to false positive BiFC signals.

Adding to these difficulties, the study of organellar PPIs by BiFC and other techniques is also hindered by the presence of targeting peptides. The position of the FP fragment fusion at the N or C terminus of a POI can affect the ability of the POI to interact with partner proteins (Miller et al., 2015). Therefore, it is common to test all FP fusions orientations in all possible combinations with the target protein to maximize the chances of identifying a PPI. However, transit peptides complicate the cloning process for FP fusions. It is likely for this reason that, to our knowledge, all BiFC experiments carried out in the chloroplasts and mitochondria to date involve POIs with C-terminus FP fragment fusions.

To further explore and overcome the challenges associated with performing organellar BiFC we developed a modular approach based on Modular Cloning (MoClo)(Weber et al., 2011; Engler et al., 2014; Gantner et al., 2018). The modular nature of the approach simplifies cloning of chloroplast proteins, and allows the assembly of all transcriptional units including a reference protein on a single construct. Using synthetic biology principles, we designed, built and tested components to arrive at an optimised system suitable for testing protein-protein interactions in the chloroplast by quantitative BiFC. Development of the system highlighted a number of factors critical for performing successful chloroplastic BiFC including the selection of negative controls, the nature of the FP split used, the cell type analysed, and the choice of reference protein for quantification. Finally, we discuss how the MoClo-based BiFC approach could be extended to other organelles and adapted to other interaction methods.

## Results

### A modular system for testing interactions by BiFC

The MoClo cloning system provides an attractive framework for the assembly and in planta expression of chimeric proteins. Using the MoClo syntax we conceived an approach for the systematic assembly of gene fusions for BiFC that we named MoClo-based BiFC or MoBiFC (Fig.1). In MoBiFC the coding sequence (CDS) of an FP fragment is fused upstream or downstream to the CDS of a protein of interest. A generic targeting presequence (e.g. encoding a chloroplast transit peptide) can be included. A second synthetic gene encoding the interaction partner to be tested is assembled in the same way with the complementary FP fragment, and the two genes are then assembled in a multigenic T-DNA along with a gene encoding the suppressor of silencing P19 (Scholthof, 2006) and a fluorescent marker of transformation (Fig. 1C).

**Fig. 1.**
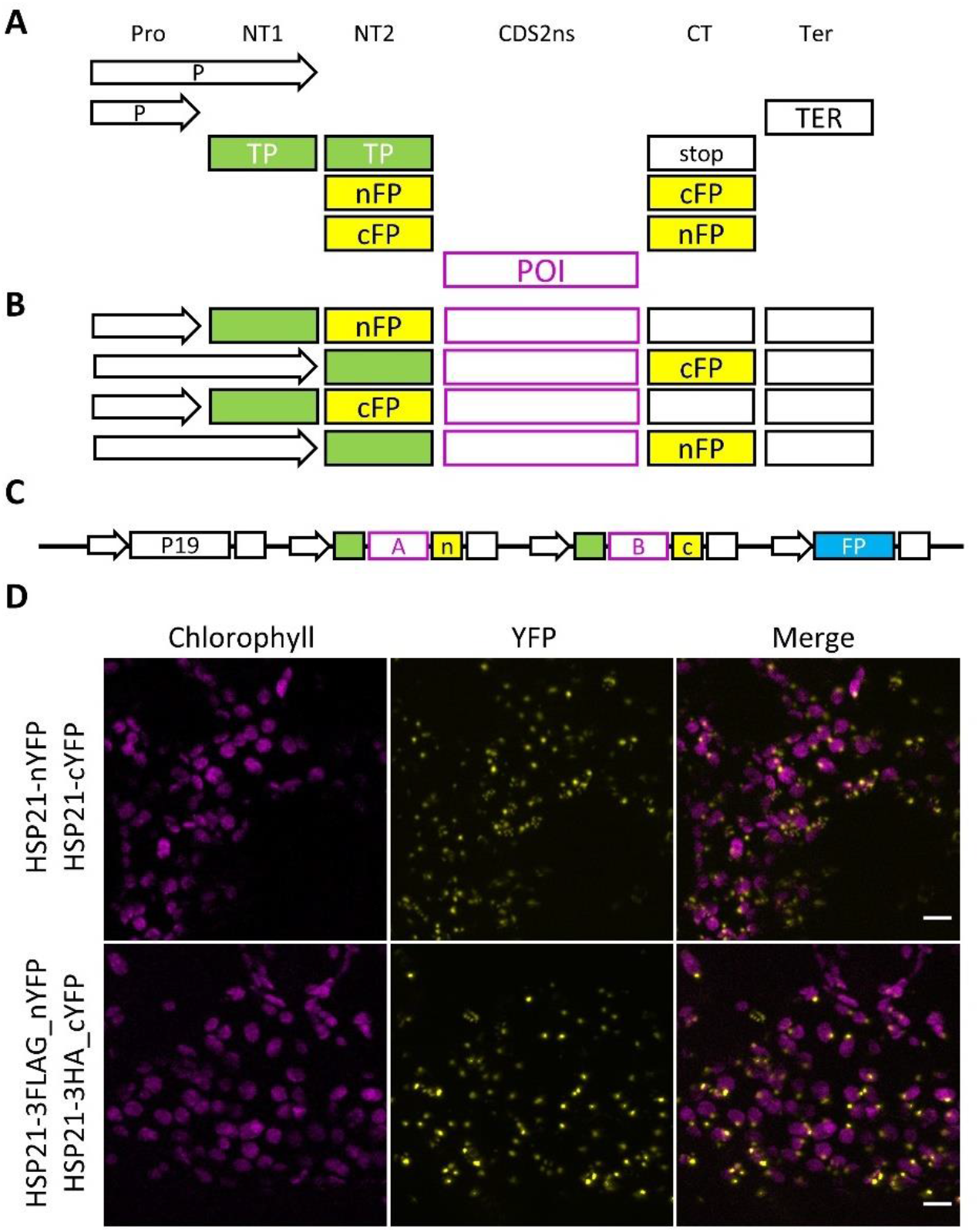
A MoClo-based approach for testing protein protein interactions by BiFC. An overview of a modular cloning-based approach for assembling (a) Level 0 components into (b) transcriptional units and then (c) multigenic assemblies for BiFC assays. The MoClo syntax of Engler et al. (2014) is indicated on the top line. P, promoter; Ter, terminator; TP, Rubisco transit peptide; FP, fluorescent protein; POI, protein of interest. (d) BiFC assay in *N. benthamiana* mesophyll cells showing the TP chloroplast targeted HSP21 self-interaction using untagged YFP fragments (nYFP/cYFP) or epitope-tagged YFP fragments (3FLAG_nYFP, 3HA_CYFP). Scale, 10 μm.

As proof of principle we tested the chloroplast protein HSP21 that is known to interact with itself (Lambert et al., 2011; Zhong et al., 2013; Rutsdottir et al., 2017). We assembled synthetic fusion genes using the mature HSP21 CDS, the two fragments of the 174/175 YFP split (Citovsky et al., 2006) and the Rubisco small subunit transit peptide (TP) for chloroplast targeting (Lee et al., 2002). YFP fluorescence was observed in the chloroplasts of *Nicotiana benthamiana* leaf mesophyll cells, indicating re-assembly of YFP due to the interaction between HSP21-nYFP and HSP21-cYFP (Fig. 1D). The signal showed the same pattern as HSP21-GFP (Fig. S1A), and was similar to the BiFC pattern reported previously for HSP21 (Zhong et al., 2013). Next, to enable detection of fusion proteins by immunoblot, we added a triple FLAG tag to the N-terminal fragment of YFP and a triple HA tag to the C-terminal fragment. When tested with HSP21 the tagged and untagged versions of the YFP fragments displayed a comparable interaction signal in terms of fluorescence intensity and localisation, indicating that the presence of the epitope tags does not disturb the BiFC (Fig. 1D). The epitope tags were also easily detectable by immunoblotting (Fig. S1B). We therefore demonstrate that the new MoClo components are functional, and that the MoClo system is suitable for assembling gene fusions for performing chloroplastic BiFC.

### MoBiFC can be used to robustly test protein-protein interactions in the chloroplast

Next, we tested the BiFC system using different partner proteins and controls. We first selected the pair HSP21 and PTAC5 that were previously shown to interact by BiFC and pull-down assays (Zhong et al., 2013). We found that expression of HSP21-nYFP and PTAC5-cYFP resulted in reconstitution of YFP fluorescence in a similar pattern to HSP21-nYFP / HSP21-cYFP in the mesophyll cells (Fig. 2). The nucleo-cytoplasmic CFP served as a useful transformation control, allowing the identification of cells containing the multigenic T-DNA. Zhong et al. (2013) showed that deletion of a C-terminal region (253-387) of PTAC5 abolished the HSP21-PTAC5 interaction. We therefore tested the interaction of HSP21 with PTAC5 lacking this C-terminal region (ΔPTAC5) to determine whether it acted as a suitable negative control. We observed a significant drop in fluorescence intensity compared to HSP21-nYFP/ PTAC5-cYFP, although we were still able to detect YFP reconstitution in a similar speckled pattern (Fig. 2, S2). While our results confirm ΔPTAC5 as a suitable negative control for PTAC5, no speckles were reported for HSP21-nYFP / ΔPTAC5-cYFP in the previous study (Zhong et al., 2013). This may be due to differences in experimental setup *(N. benthamiana* mesophyll cells here versus Arabidopsis protoplasts previously) or post acquisition analysis.

**Fig. 2.**
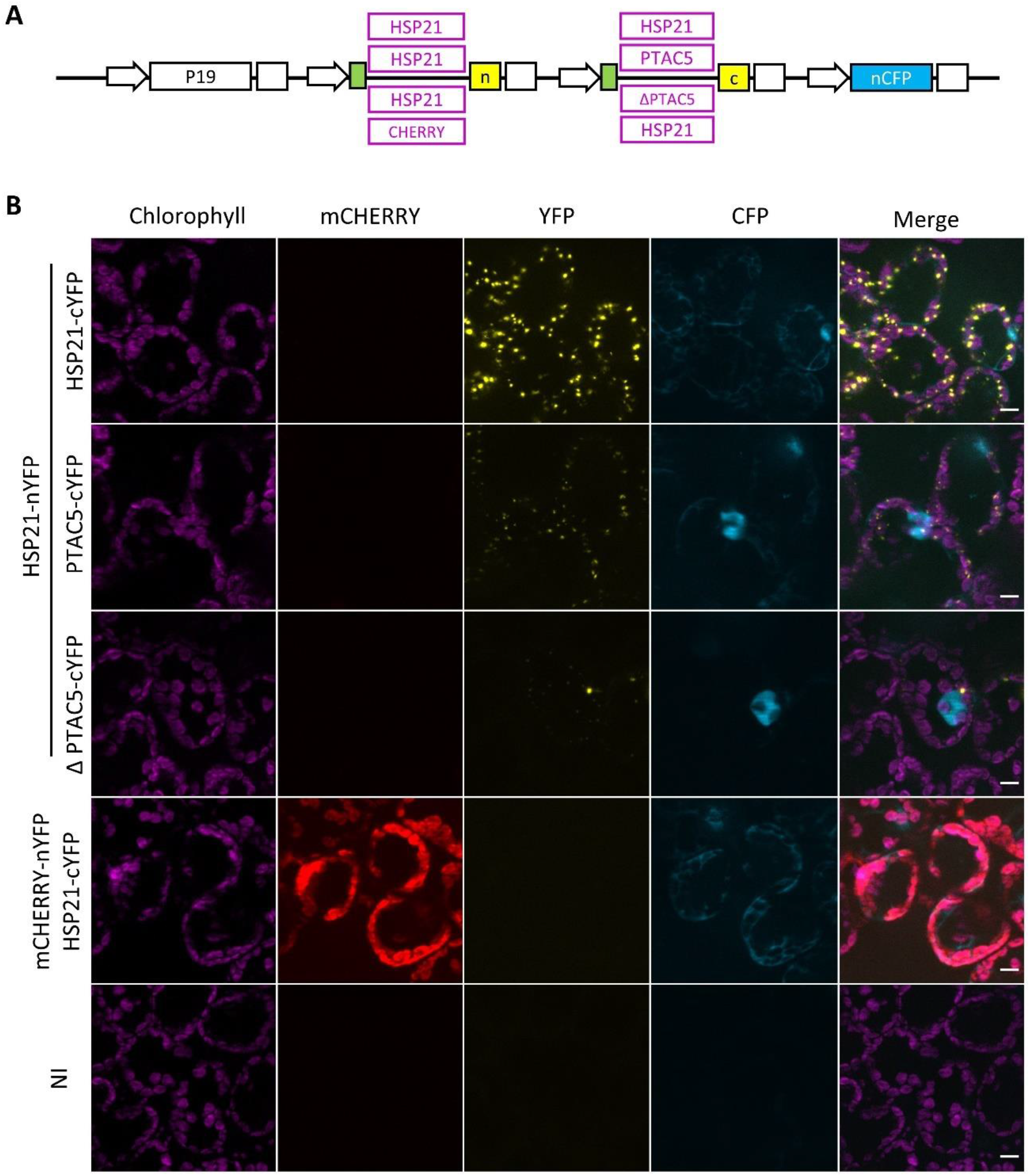
mCHERRY-nYFP and HSP21-cYFP do not produce a non-specific assembly signal in the chloroplast. (a) The POI region of the multigenic BiFC construction was varied to produce four different chloroplast targeted HSP21 pairs that were used for (b) BiFC assays in *N. benthamiana* mesophyll cells. NI, not inoculated; scale, 10 μm.

We next searched for a negative control with the potential to be used in a more general fashion. We selected the monomeric fluorescent protein mCHERRY as a candidate (Shaner et al., 2004). Chloroplast-targeted mCHERRY-nYFP showed strong mCHERRY fluorescence in the chloroplasts, and a very low BiFC signal in the presence of HSP21-cYFP expressed from the same T-DNA. The mCHERRY BiFC signal showed no speckles, and was significantly lower than the HSP21-nYFP / HSP21-cYFP signal (Fig. S2). Surprisingly, mCHERRY-nYFP / HSP21-cYFP showed a higher signal than HSP21-nYFP / ΔPTAC-cYFP, despite the presence of visible speckles for the second pair. These differences in signal are likely due to differences in protein localisation and abundance, and highlight the importance of these factors in making comparisons between BiFC pairs. Together, these results indicate that mCHERRY can act as a useful generic negative control for BiFC. mCHERRY has the advantage of showing a homogenous stroma localisation, and fluorescence that can be visualised directly during microscopy to allow rapid assessment of BiFC interaction specificity.

### MoBiFC allows N-terminal fusion proteins to be tested within the chloroplast

The MoBiFC cloning approach allows the introduction of an N-terminal tag while preserving organellar targeting presequences (Fig. 1). To test whether BiFC is still functional in this configuration we performed BiFC assays with the YFP fragments fused to the N-terminal of the proteins of interest (Fig. 3). N-terminal tagged HSP21 showed a similar BiFC signal to C-terminal tagged HSP21, indicating that chloroplast import, HSP21/HSP21 interaction and YFP reconstitution are not hindered. At the same time, the background signal for the mCHERRY negative control was slightly lower for the N-terminal nYFP fusion than the C-terminal nYFP fusion. Therefore, the MoClo based BiFC approach allows the straightforward generation of N- and C-terminal tagged protein fusions for chloroplastic BiFC assays.

**Fig. 3.**
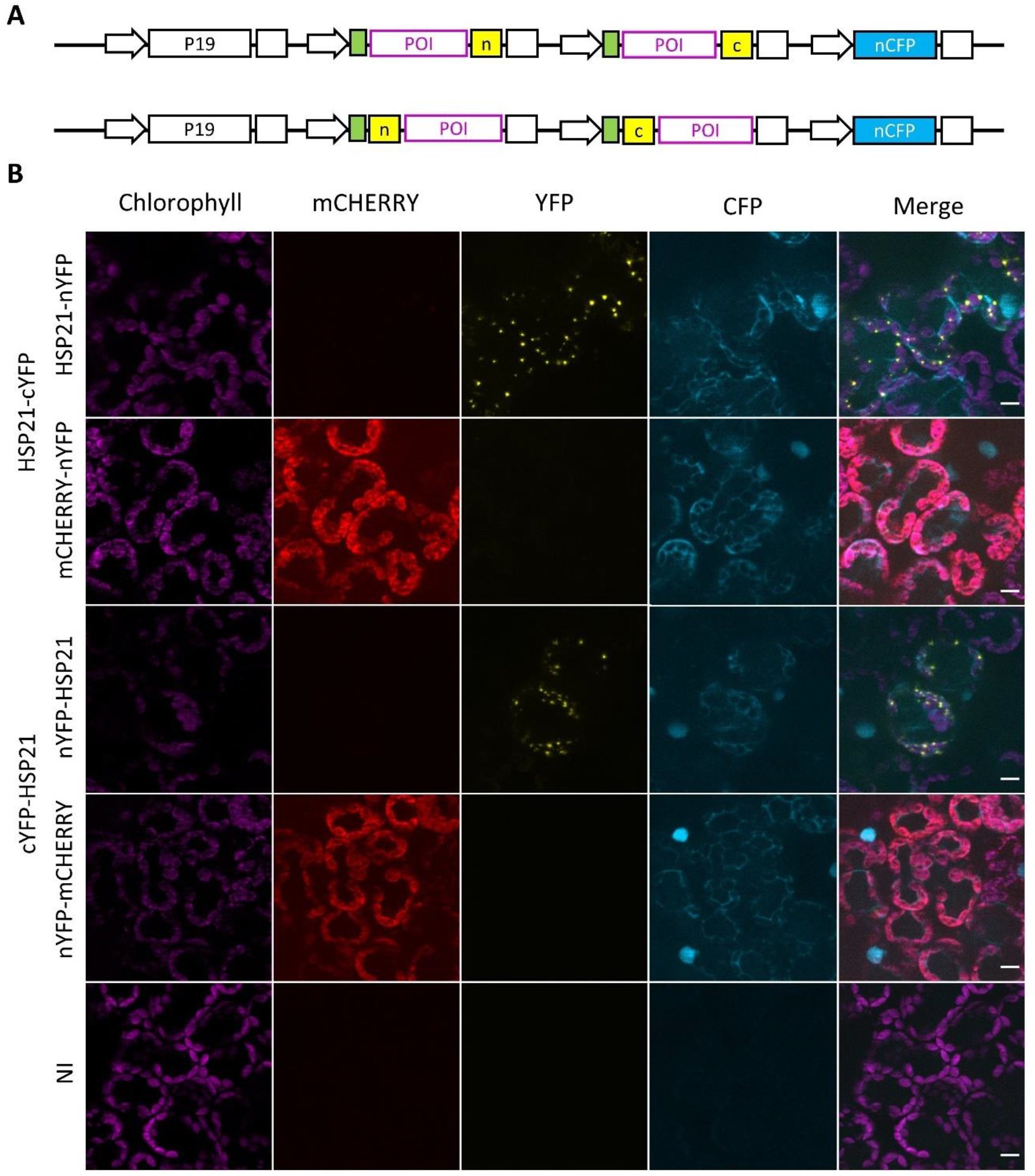
N-terminal FP fusions function similarly to C-terminal fusions in BiFC experiments. (a) Chloroplast targeted HSP21 and HSP21 / mCHERRY pairs with C-terminal and N-terminal FP fragment fusions were used for (b) BiFC assays in *N. benthamiana* mesophyll cells. NI, not inoculated; scale, 10 μm.

### Optimisation of MoBiFC with different promoters, FP splits and reference FPs

The modular nature of the MoClo system allows different components to be swapped in and tested. We therefore sought to determine whether altering the fluorescent protein split or promoters could improve performance using the HSP21 pair (Fig. 4). We first replaced the 174/175 YFP split with the 210/211 mVENUS split that was reported to display a lower level of non-specific assembly (Gookin and Assmann, 2014). The HSP21-nVENUS / HSP21-cVENUS interaction pattern was very similar to the YFP split, while the strength of the fluorescence signal doubled and nVENUS fusion protein accumulation also increased (Fig. 4, S3). However, the mCHERRY-nVENUS / HSP21-cVENUS negative control produced almost 4 times higher non-specific signal than for the corresponding YFP split (Fig. S3). A possible explanation for the higher non-specific signal is self-complementation within the mCHERRY-nVENUS fusion protein, or fluorescence bleed-through from mCHERRY into the YFP channel. However, we found that the non-specific signal was detectable only when mCHERRY-nVENUS was expressed with HSP21-cVENUS (Fig. S4). Therefore, the mVENUS split increases the BiFC signal intensity, but this can be accompanied by an increase in non-specific assembly in at least some cases.

**Fig. 4.**
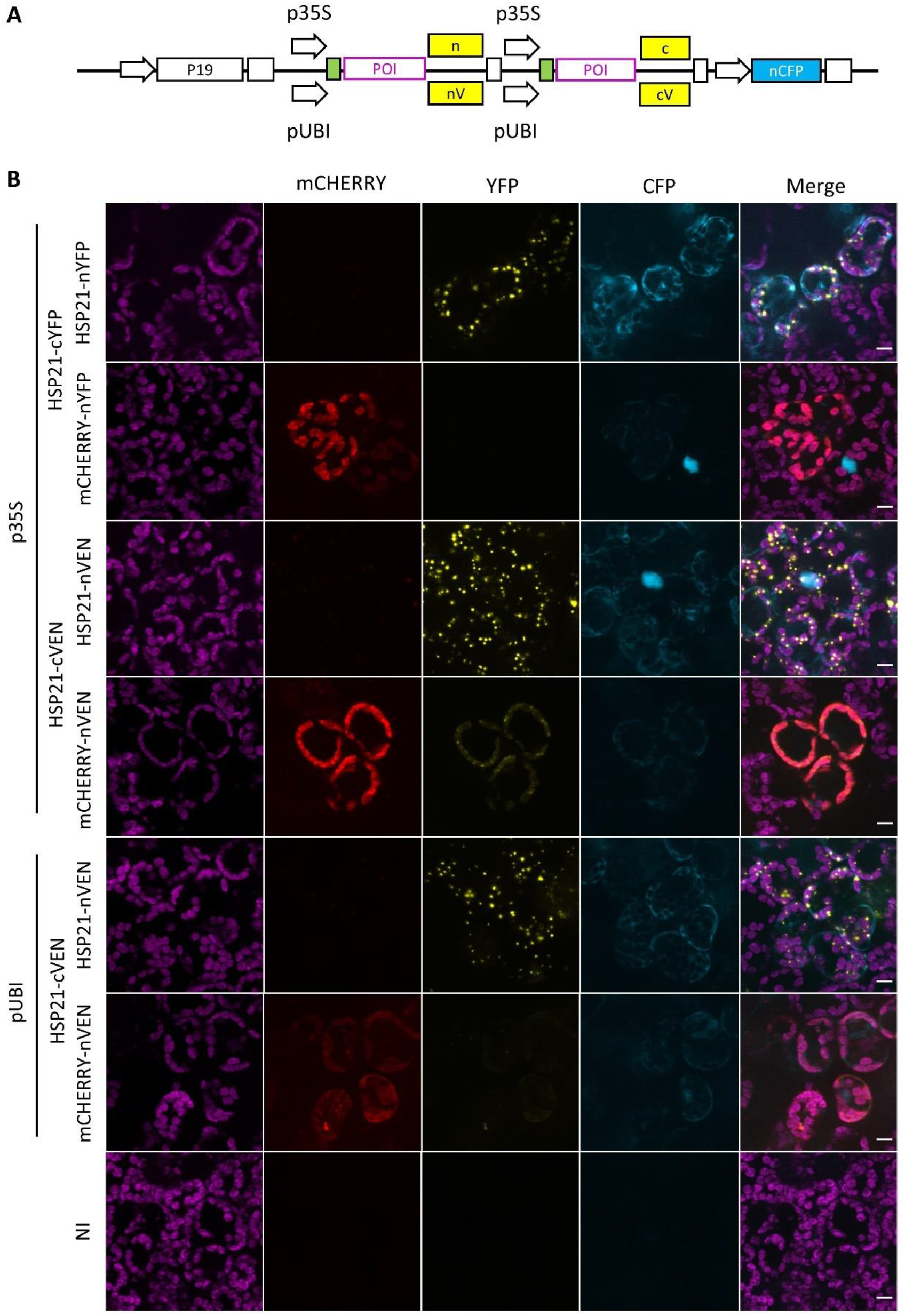
The sensitivity of the BiFC can be adjusted with different FP splits and promoters. (a) The promoter and FP fragment regions in the multigenic BiFC construct were varied to produce six different chloroplast targeted HSP21 pairs that were used for (b) BiFC assays in *N. benthamiana* mesophyll cells. NI, not inoculated; nV/cV, mVENUS fragments; nVEN/cVEN, mVENUS fragments. Scale, 10 μm.

The strong 35S promoter of the cauliflower mosaic virus was used to drive expression of the fusion genes in the previous experiments. High expression levels could be responsible for driving non-specific assembly of the mVENUS split. We therefore attempted to reduce expression levels to improve the signal to noise ratio. For this purpose, we used the Arabidopsis *UBQ10* promoter which was shown to be substantially weaker than the 35S promoter in tobacco (Grefen et al., 2010). While the fluorescence intensity was somewhat reduced for the HSP21 / HSP21 pair we still observed the diffuse mVENUS signal in the mCHERRY / HSP21 negative control (Fig. 4, S3). Along with the reduction in fluorescence, immunoblotting showed a small but clear decrease in levels of fusion proteins produced using the *UBQ10* promoter (Fig. S3). Taken together, our attempts at optimisation indicate that the 35S promoter and the 174/175 YFP split modules are currently the best choice for BiFC in the chloroplast under these conditions.

We next turned to the reference FP. While useful, the nucleo-cytosolic CFP shows a rather weak fluorescence and is sometimes difficult to find in mesophyll cells. The signal is also not proportional to chloroplast area within a region of interest, so is not suitable for ratiometric BiFC quantification despite its presence on the same T-DNA. We therefore swapped out the nucleo-cytosolic CFP from the BiFC T-DNA and replaced it with either chloroplast targeted CFP (TP-CFP) or an OEP7 monomeric Turquoise (mTRQ) fusion (OEP7-mTRQ) (Fig. 5). OEP7 is a protein of the chloroplast outer envelope membrane, and OEP7-GFP was previously used as an outer envelope membrane marker (Lee et al., 2001). mTRQ is a monomeric variant of CFP with a higher quantum yield (Goedhart et al., 2012; Hecker et al., 2015; Gantner et al., 2018). Both TP-CFP and OEP7-mTRQ clearly marked the chloroplasts in transformed cells of the epidermis and mesophyll and showed a more intense fluorescence than the nucleo-cytosolic CFP. The signal for the HSP21-nYFP / HSP21-cYFP interaction in the chloroplasts was also preserved. However, cells transformed with TP-CFP regularly showed mCHERRY and YFP signals in the cytosol (Fig. S5). This indicates that co-expression of TP-CFP interferes with the chloroplast localisation of HSP21 and mCHERRY chloroplast. The integrity of the chloroplasts did not appear to be affected, suggesting that the mis-localisation may therefore be linked to saturation of the chloroplast import machinery. In contrast, we never observed cytosolic mCHERRY or YFP signals in mesophyll cells co-expressing OEP7-mTRQ. OEP7 insertion in the chloroplast membrane does not compete with Rubisco SSU import and is independent of known chloroplast translocons (Salomon et al., 1990; Schleiff et al., 2001). Therefore, it is likely that the use of OEP7-mTRQ prevents the saturation of the chloroplast import machinery that appears to occur when multiple chloroplast targeted proteins are co-expressed at high levels. Importantly, we also found that OEP7-mTRQ fluorescence was proportional to mCHERRY-nYFP fluorescence, which varies between cells. The proportional accumulation of OEP7-mTRQ combined with its neutral chloroplast localisation therefore allows accurate ratiometric quantification of specific and non-specific chloroplast based BiFC signals (Fig. 5).

**Fig. 5.**
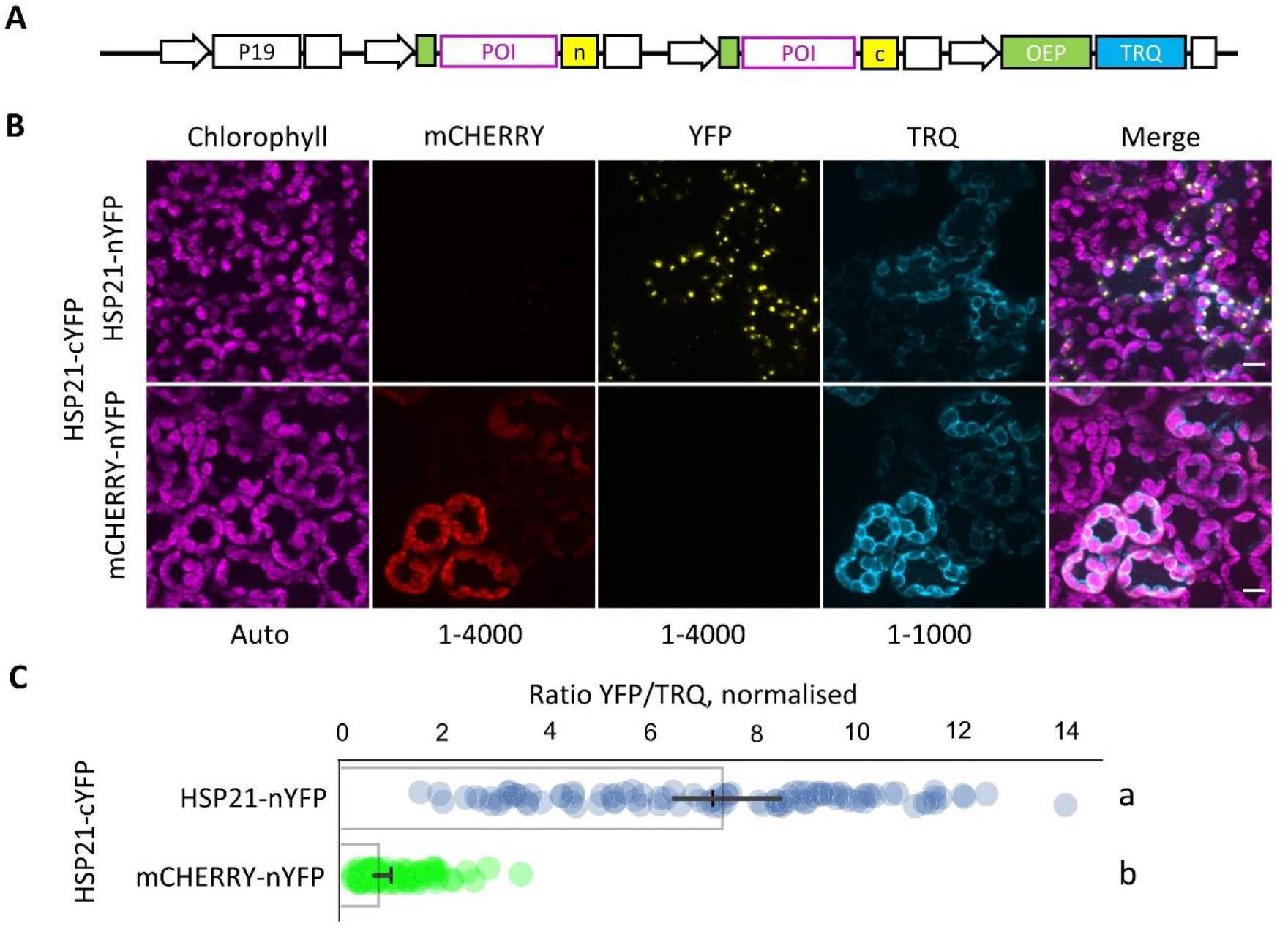
OEP7-mTRQ is a suitable reference FP for ratiometric chloroplastic BiFC. (a) Outline of multigenic BiFC construct encoding an OEP7-mTRQ reference FP that was used for (b) BiFC assays between two protein pairs in *N. benthamiana* mesophyll cells. Scale, 10 μm. (c) Normalised BiFC signals were calculated as the ratio between total eYFP and TRQ fluorescence and normalised to HSP21-cYFP / mCHERRY-nYFP. Bar indicates mean and cross indicates median +/− 95% confidence interval (n=100 transformed cells). Significance was calculated using the Kruskal Wallis test, groups are indicated by lower case letters (*P*<0.001).

### Non-specific assembly in different *N. benthamiana* cell types

During the numerous experiments carried out in *N. benthamiana,* we noticed that different tissues did not show the same response in the BiFC assays. Mesophyll cells from the upper and lower side of the leaf always showed comparable results. However, epidermal cells often showed BiFC signals for the HSP21-nYFP / HSP21-cYFP pair that were much stronger and differently distributed than those observed in the mesophyll (Fig. S6). Furthermore, the negative control mCHERRY-nYFP / HSP21-cYFP showed a non-specific BiFC signal. This non-specific assembly occurred even after short incubation times following infiltration (24-48 h), using low Agrobacterium infiltration concentrations, and with the mVENUS split. Interestingly, BiFC assays in Arabidopsis epidermal cells using the same constructions showed a specific and correctly localised HSP21-nYFP /HSP21-cYFP interaction without non-specific mCHERRY-nYFP / HSP21-cYFP assembly (Fig. S7). However, the signal intensity in Arabidopsis was considerably lower than in *N. benthamiana* epidermal cells. Taken together, these observations might therefore suggest that the combination of high expression levels in *N. benthamiana* and the small size of epidermal chloroplasts can lead to protein overaccumulation within the chloroplasts. High protein concentrations will favour the spontaneous self-assembly of the YFP split and will also increase the formation of protein aggregates with incorrect localisation.

### MoBiFC modules can be rapidly adapted for other approaches

The universal MoClo assembly syntax allows the re-use of Level 0 modules for other applications. BiFC is mostly performed using two known POIs, and candidate POIs must first be identified using other methods such as co-immunoprecipitation followed by mass spectrometry (MS). PPI identification has recently received a boost thanks to new proximity labelling approaches such as the use of the optimised promiscuous biotin ligase TurboID that, when fused to a POI, biotinylates proteins within the immediate vicinity (Branon et al., 2018). Proteins biotinylated by the POI-TurboID fusion can then be isolated and identified by MS. Several studies now show that TurboID functions well for PPI identification in plants, and is even able to identify transient interactions such as those between a kinase and its substrate (Kim et al., 2019; Mair et al., 2019; Zhang et al., 2019; Arora et al., 2020). However, to our knowledge TurboID or any other proximity labelling method have not yet been demonstrated in the chloroplast. We therefore designed TurboID modules in the MoClo syntax for N and C-ter fusions to chloroplast targeted POIs (Fig. S8, Tables S1). A chloroplast targeted YFP-TurboID fusion localises correctly to the chloroplast where it biotinylates a range of proteins including a co-expressed chloroplast targeted CFP (Fig. S8). While this experiment shows that TID is functional within the chloroplast, it also highlights the promiscuous nature of TID. For the identification of genuine interactors a non-interacting TID control will therefore be required allow the quantitative demonstration that a POI preferentially labels a specific subset of chloroplast proteins. This addition to the toolkit will complement the BiFC tools by facilitating the de novo identification of POIs within the chloroplast and other cellular compartments.

## Discussion

We describe the development of a new BiFC approach and toolkit for the investigation of protein-protein interactions in the chloroplast. The approach facilitates the cloning process for chloroplast-targeted proteins, allows robust ratiometric quantification, and the toolkit comes with model positive and negative controls for chloroplastic BiFC (Table S1). Furthermore, the open design of MoClo based systems facilitates user-driven optimisation and enrichment with new modules and functions. As an example, we demonstrate a complementary TurboID extension for identification of candidate interactors by proximity labelling. Our study also highlighted the pitfalls that can occur in setting up chloroplastic BiFC assays. We find that the choice of FP split, negative controls, cell type, and even reference FP localisation can have major effects on the outcome of a BiFC experiment.

We show that chloroplast targeted mCHERRY is a useful generic negative control for chloroplast BiFC assays due to its intrinsic fluorescence and homogenous stromal localisation (Fig. 2-5). A similar approach was used previously for a nuclear BiFC assay where the related TagRFP was used as negative control for BiFC in the nucleus (Horstman et al., 2014). Here, we show that the FP fragment fusion to mCHERRY does not restore YFP/mVENUS fluorescence on its own (Fig. S4). Furthermore, the BiFC signal we observed from the nonspecific assembly of mCHERRY-nYFP and HSP21-cYFP due to high or mislocalised expression (Fig. 5, Fig. S6,) indicates that fusion of YFP fragments to mCHERRY does not compromise YFP re-assembly. While mCHERRY is a useful tool, we do however recommend that, where possible, a mutated version of the POI or a protein closely related to the POI is also used as a negative control because these proteins are more likely to share the same localisation and abundance as the POI (Kodama and Hu, 2012; Gookin and Assmann, 2014; Horstman et al., 2014; Kudla and Bock, 2016). Specific interactions may also be demonstrated by the expression of a non-labelled version of the POI that competes with the FP fragment labelled version to deplete the BiFC signal (Kodama and Hu, 2012; Bischof et al., 2018). MoClo cloning would facilitate competition assays by allowing the straightforward addition of a transcriptional unit for an unlabelled competitor protein to the multigenic BiFC expression construct.

We found a better signal-to-noise ratio with the 174/175 YFP split (Citovsky et al., 2006) than with the 210/211 mVENUS split (Gookin and Assmann, 2014). This was surprising because the mVENUS split was reported to display a particularly low level of non-specific assembly. The better performance of the 174/175 YFP split may be due to the environment of the chloroplast stroma, the specific proteins tested here, or the nature of the expression system itself. These results suggest that it may be advantageous to test more than one FP split for BiFC depending on the cellular compartment or even protein pair tested.

We investigated model stromal protein interactions in this study. In the future it would be interesting to investigate how the approach might be extended to other chloroplast compartments and to other organelles. Indeed, BiFC experiments have shown interactions between stromal proteins and integral thylakoid membrane proteins with C-terminal FP fusions (Wang and Blumwald, 2014; Ouyang et al., 2020). MoBiFC has the potential to be simply extended to other organelles such as the mitochondria, peroxisome and nucleus by the replacement of the SSU CTP modules with specific organellar targeting modules. However, care would need to be taken to avoid saturation of the import machinery and overaccumulation of the fusion proteins within the organelle as we observed here in the chloroplast under specific circumstances (Fig. 5, Fig. S6). Due to its open design and the ease with which multigenic transformation constructs can be made the MoBiFC toolkit is also extendable to multicolour BiFC, the inclusion of competitor controls as discussed above, systematic library screening, and emerging techniques such as proximity labelling.

## Acknowledgements

This work was supported by Agence Nationale de la Recherche funding (G4PLAST, ANR-17-CE13-0005). MM was supported by an Erasmus international mobility grant and a mobility grant of the University of Tunis El Manar. Fluoresence microscopy experiments were performed on the PiCSL-FBI core facility, member of the France-BioImaging national research infrastructure (ANR-10-INBS-04). We thank Pascaline Auroy-Tarrago for facilitating the development of synthetic biology initiatives within the BIAM.

## Contributions

F.V., M.S., S.D., C.L., P.C and B.F. conceived and planned the experiments. F.V., M.S., M.M., S.D., P.C., and B.F. performed the experiments. F.V., M.M., S.D., C.L., P.C. and B.F contributed to the analysis and interpretation of the results. F.V., C.L., P.C and B.F. wrote the manuscript. All authors provided critical feedback and helped shape the research, analysis and manuscript.

## Materials and Methods

### Plant material and growth conditions

*Nicotiana benthamiana* was used for all experiments in this study unless otherwise indicated. Arabidopsis plants carrying the GVG-AvrPto transgene were used for Arabidopsis BiFC assays (Tsuda et al., 2012). Both Arabidopsis and *Nicotiana* plants were grown in a controlled environment at 120 μmol m^-2^ s^-1^ illumination with an 8 h / 16 h photoperiod at 22°C day / 20°C night, and 55% day / 75% night relative humidity.

### Cloning

New gene parts (level 0 modules) were amplified by PCR or synthesised directly (Twist Biosciences) and sequenced. The resulting modules are free from BsaI, BsmBI, BpiI and SapI Type IIS sites and can be mobilised in the MoClo, Goldenbraid and Loop cloning systems (Sarrion-Perdigones et al., 2013; Patron et al., 2015; Pollak et al., 2019). Restriction ligation reactions for the assembly of transcriptional units (Level 1) and assemblies of transcriptional units (Level 2) were performed using the single step protocol described previously with small modifications (Weber et al., 2011) and according to the MoClo Golden Gate assembly standard (Engler et al., 2014). Briefly, 100 fmol of each insert plasmid and 50 fmol of acceptor plasmid were mixed with restriction enzyme (BpiI or BsaI) and T4 DNA ligase in T4 ligase buffer in 20 μl reactions and incubated at 37 °C for 5 h. A 1.5 μl aliquot was transformed into DH10B cells by electroporation and transformants selected on an appropriate antibiotic. Correct assembly was confirmed by digestion. Lists of new modules created and their sequences are provided in the Table S1, Table S2 and Supplementary file 1. All other Level 0 and infrastructure modules used were described previously (Engler et al., 2014; Gantner et al., 2018). The vectors containing the principal modules for the toolkit have been deposited at Addgene (Table S1).

### Transient expression by agroinfiltration

*Agrobacterium tumefaciens* GV3101 transformed with Level 1 transcriptional units or Levels 2 multigenic units were grown at 28°C overnight in LB medium supplemented with rifampicin and a selective antibiotic. The cultures were then diluted to an OD600 of 0.1 in infiltration buffer containing 10 mM MES pH 5.5, 10 mM MgCl2 and 200 μM acetosyringone were infiltrated into leaves of one month old *N. benthamiana* plants using a 1 ml syringe. For Arabidopsis transient expression, leaves of GVG-AvrPto plants were sprayed with 2 μM dexamethasone 24 h before inoculation (Tsuda et al., 2012), and then infiltrated as described for *N. benthamiana.* Infiltrated plants were returned to standard growth conditions for three days before observation.

### Bimolecular fluorescence complementation assays

All BiFC assays were repeated at least twice and showed similar results. Leaf discs were taken 3 days after agroinfiltration. The discs were mounted in perfluorodecalin to allow visualisation of mesophyll cells as described previously (Littlejohn and Love, 2012). Capture of fluorescence images was performed using the AxioImager APO Z1 microscope (Zeiss). The following filters were used: chlorophyll, excitation 625-655 nm, emission 665-715 nm; mCHERRY, excitation 533-558 nm, emission 570-640 nm; eYFP/mVENUS, excitation 490-510 nm, emission 520-550 nm; and CFP/mTRQ, excitation 431-441 nm, emission 460-500 nm. Standard exposure times of 5 ms for chlorophyll and 75 ms for FPs was kept for all observations. Images were captured from different regions of each inoculated leaf, and from at least two leaves per experiment. 10 μm deep Z stacks composed of 21 slices were acquired and then converted into maximum intensity projections. During post-acquisition processing the images of each experiment were treated identically. Histogram levels were set so that there were no oversaturated pixels, except for images in Fig S2, S3 and S4 where histogram levels were modified in the same way for all images to show low intensity signals. For quantification of normalised fluorescence intensities unprocessed images were used, and regions of interest containing only transformed cells (i.e those displaying reference CFP fluorescence) were defined in ImageJ (Schindelin et al., 2012; Schneider et al., 2012). The integrated signal density was calculated for the YFP/VENUS channel in each region of interest and divided by the total chlorophyll or CFP fluorescence in the same region to provide a fluorescence intensity ratio. Graphs and statistical tests were generated in Python (Python Software Foundation, https://www.python.org/) using the Panda (McKinney, 2010), Matplotlib (Hunter, 2007), Seaborn (Waskom et al., 2020) and Pingouin (Vallat, 2018) packages.

### Biotin labelling

Leaf discs were taken from *N. benthamiana* plants three days after agroinfiltration and infiltrated with a solution of 50 μM biotin. After incubation for one hour under plant growth conditions discs were frozen and total proteins extracted as described below.

### Immunoblotting and protein detection

Total leaf proteins were extracted in SDS sample buffer and separated by SDS-PAGE as described previously (Sugliani et al., 2016). Proteins were transferred onto a nitrocellulose membrane and probed with specific antibodies. Primary antibodies targeting the HA tag (monoclonal ab9110, Abcam), FLAG tag (monoclonal 637301, BioLegend) and GFP (polyclonal A-11122, Thermofisher) were all used at a concentration of 1/3000. Biotinylated proteins were detected directly using streptavidin-horse radish peroxidase conjugate (RPN1231, Cytiva). Total proteins were visualised after separation using SYPRO ruby protein stain (Thermofisher).

## Supplementary Data

**Fig. S1. HSP21-GFP localisation and detection of 3FLAGnYFP and 3HAcYFP**

**Fig. S2. Quantification of the BiFC experiments shown in Fig. 2.**

**Fig. S3. Related to Fig. 4, the sensitivity of the BiFC can be adjusted with different FP splits and promoters.**

**Fig. S4. mCHERRY FP fragment fusions do not show YFP fluorescence when expressed alone.**

**Fig. S5. Chloroplastic CFP is unreliable as a reference FP.**

**Fig. S6. Epidermal cells have a high rate of false positive signals.**

**Fig. S7. BiFC in Arabidopsis.**

**Fig. S8. Functional test of a chloroplast TurboID module.**

**Table S1 List of principal modules for MoBiFC (submitted to Addgene)**

**Table S2 List of all constructs generated.**

## Supplementary File 1

**Fig. S1.**
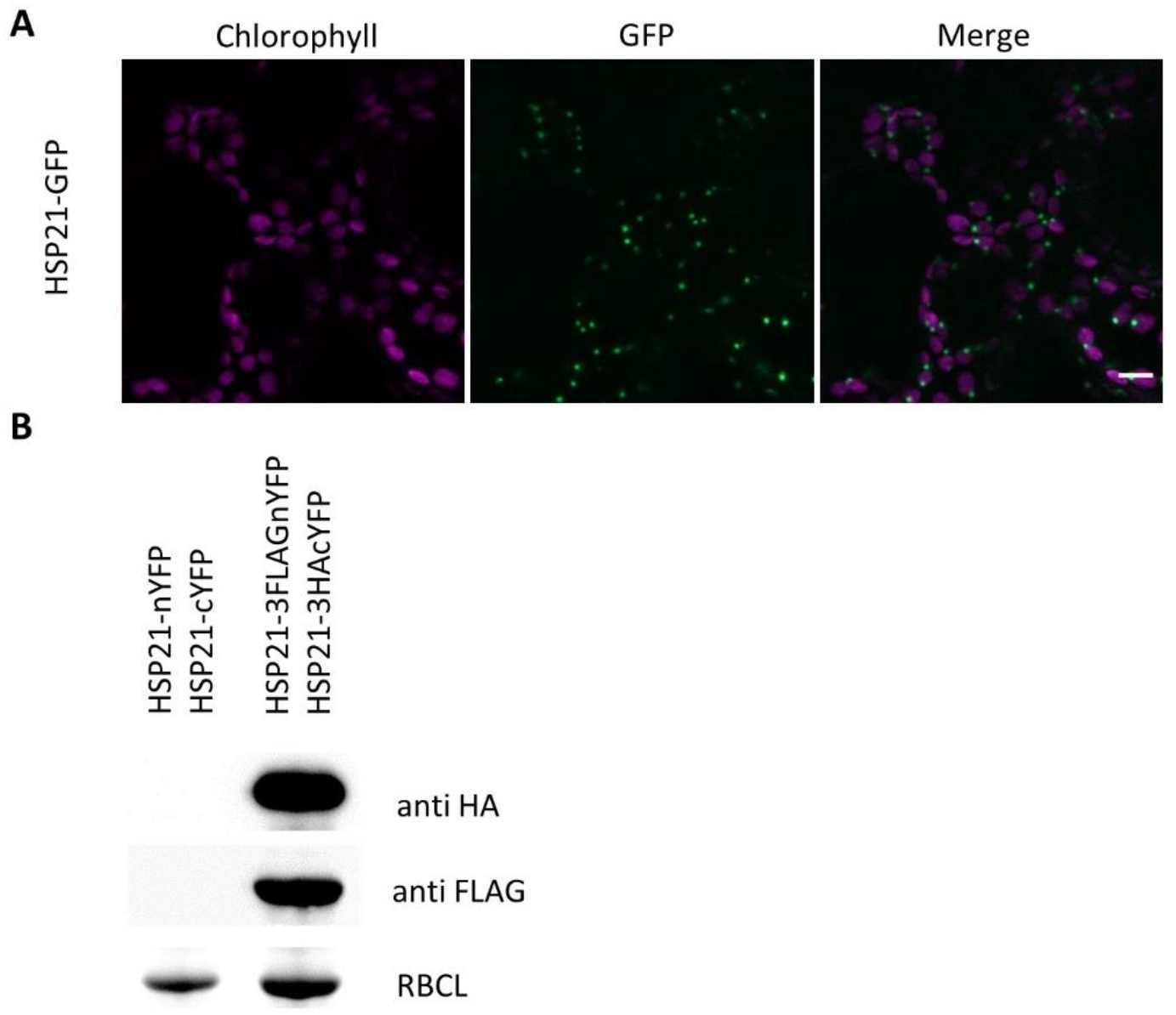
HSP21-GFP localisation and detection of 3FLAGnYFP and 3HAcYFP. (a) Localisation of HSP21-GFP in *N. benthamiana* mesophyll cells. Scale, 10 μm. Standard histogram levels are indicated below each channel. (b) Immunoblots with the indicated antibodies on protein samples normalised on a weight basis from the BiFC experiments in Fig. 1. The large subunit of Rubisco (RBCL) was visualised by Sypro fluorescent total protein stain.

**Fig. S2.**
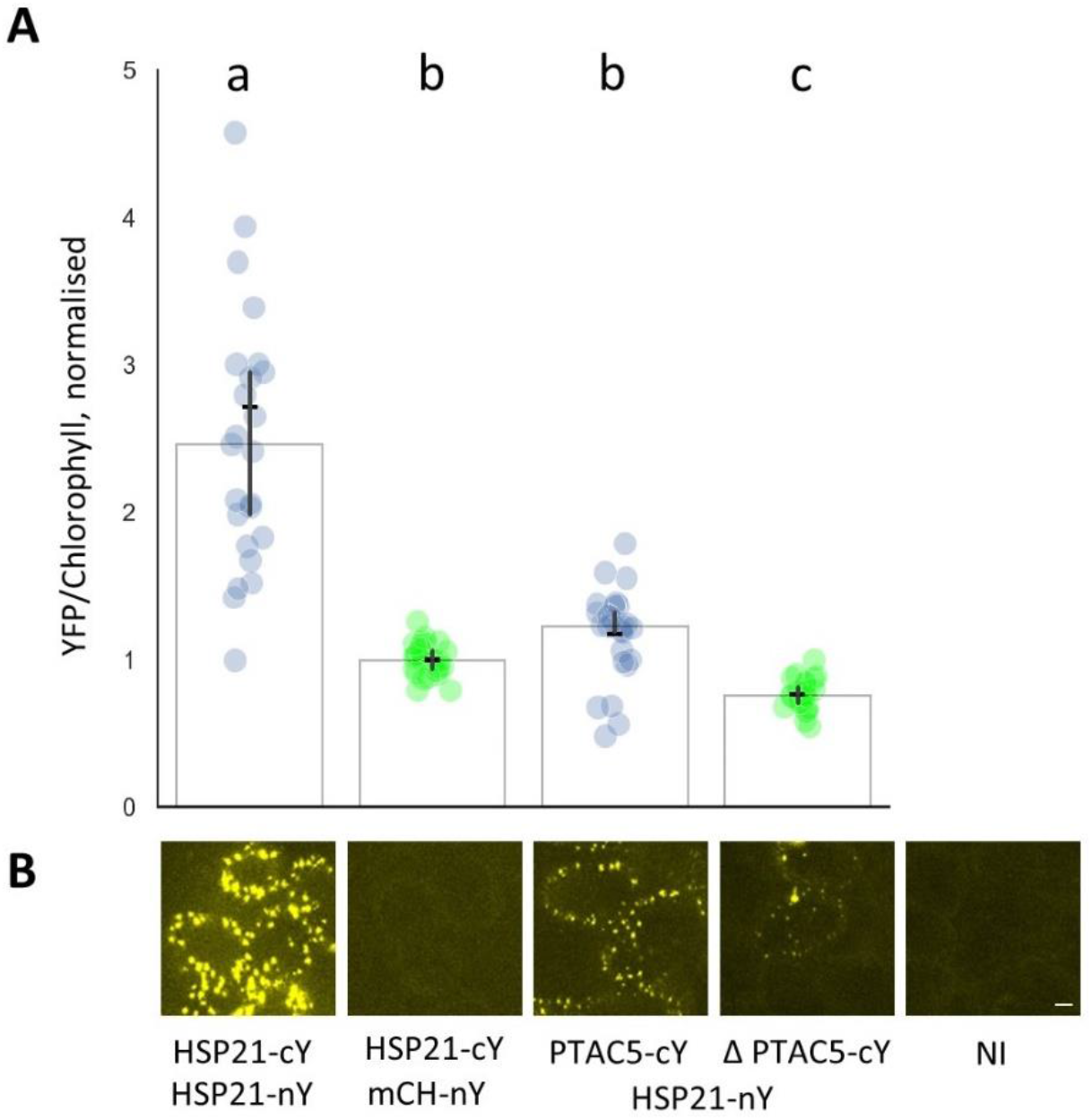
Quantification of the BiFC experiments shown in Fig. 2. (a) Normalised BiFC signals were calculated as the ratio between total eYFP/mVENUS and total chlorophyll fluorescence and normalised to HSP21-cYFP / mCHERRY-nYFP. Bar indicates mean and cross indicates median +/− 95% confidence interval (n=25-22 transformed cells). Significance was calculated using the Kruskal Wallis test, groups are indicated by lower case letters (*P*<0.001). (b) YFP channel images from Fig. 4 shown with a lower maximum histogram setting (i.e saturated) to visualise low intensity signals. nY, nYFP; cY, cYFP; mCH, mCHERRY; NI, not innoculated.

**Fig. S3.**
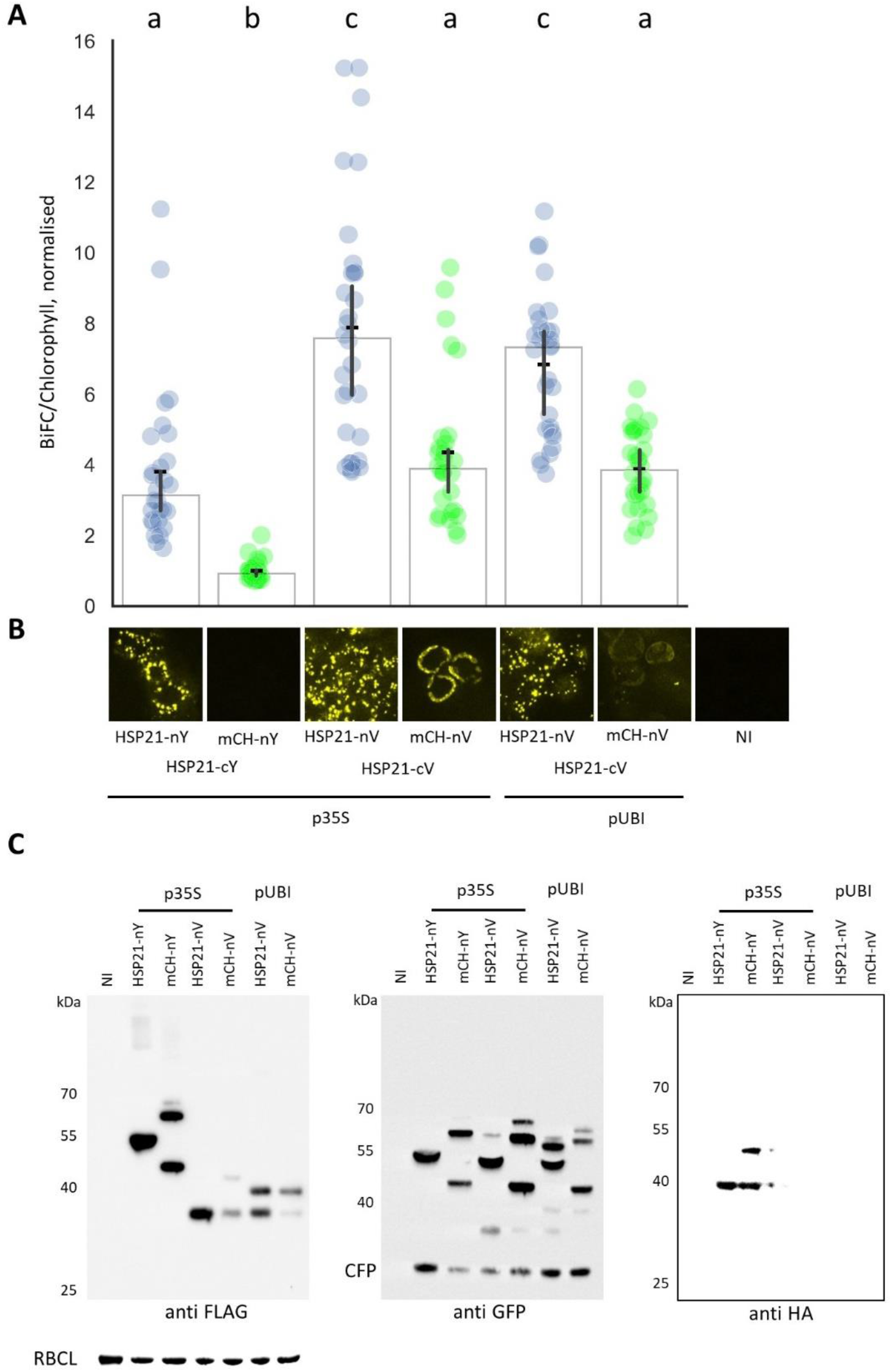
Related to Fig. 4, the sensitivity of the BiFC can be adjusted with different FP splits and promoters. (a) Quantification of the BiFC experiments shown in Fig. 4. The normalised BiFC signal was calculated as the ratio between total eYFP/mVENUS and chlorophyll fluorescence and normalised to HSP21-cYFP / mCHERRY-nYFP (arbitrarily set to 1). Bars indicate mean and crosses indicate median +/− 95% confidence interval (n=28-30 transformed cells). Significance was calculated using the Kruskal Wallis test, groups are indicated by lower case letters (*P*<0.001). (b) BiFC signal from Fig. 4 shown with a lower maximum histogram setting (i.e saturated) to visualise background signal present in HSP21-cVENUS / mCherry-nVENUS experiments. (c) Immunoblots with the indicated antibodies on protein samples normalised on a fresh-weight basis from the BiFC experiments in Fig. 4. anti-FLAG recognises the FLAG tag in 3XFLAG_nYFP (nY) and 3XFLAG_cVENUS (cV); anti-GFP recognises CFP, 3XFLAG_nYFP (nY), and nVENUS (nV); and anti-HA recognises cYFP-3XHA (cY). The large subunit of Rubisco (RBCL) was visualised by Sypro fluorescent total protein stain. mCH, mCHERRY; NI, not innoculated.

**Fig. S4.**
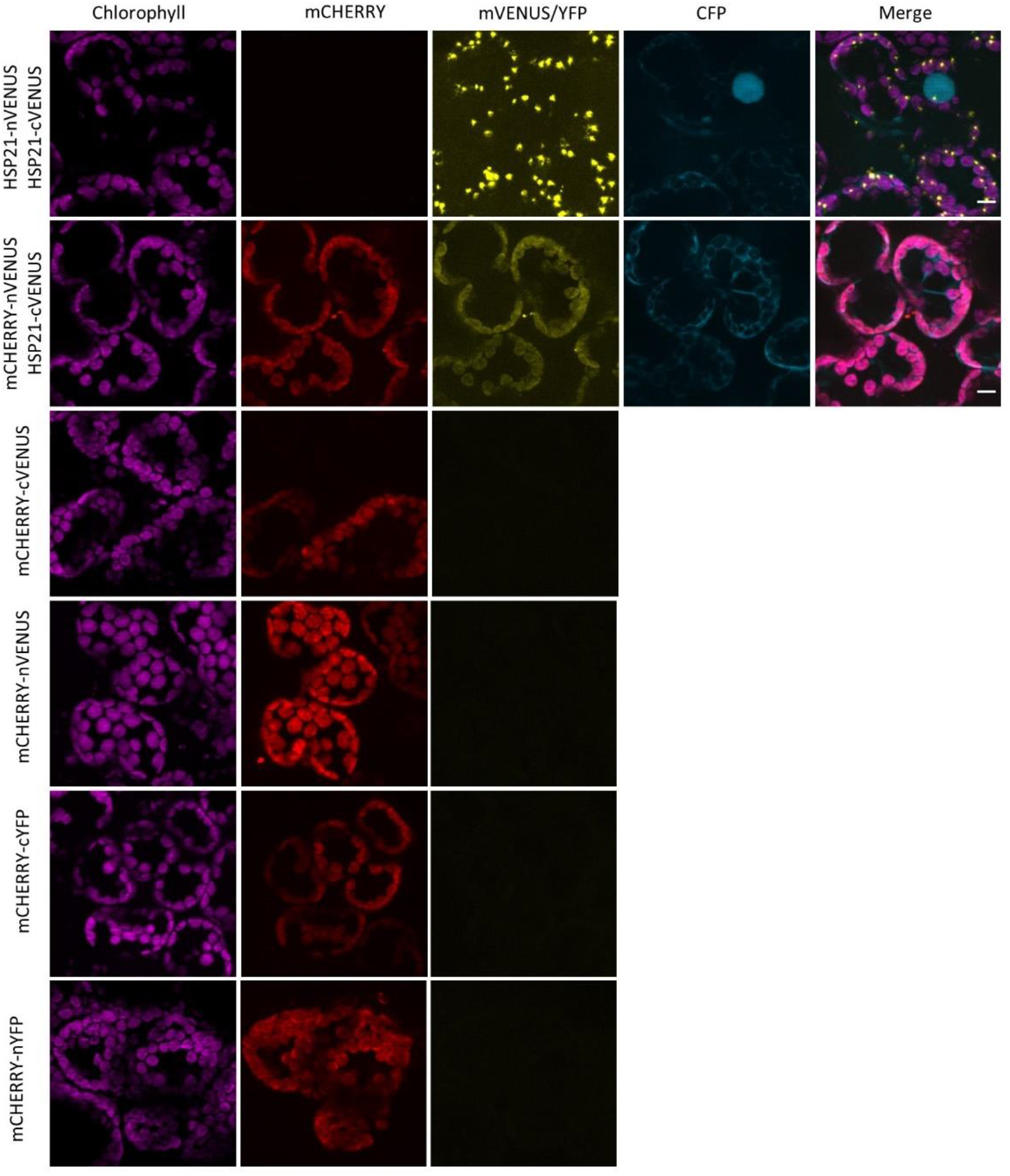
mCHERRY FP fragment fusions do not show YFP fluorescence when expressed alone. BiFC signals in *N. benthamiana* epidermal cells with the indicated proteins. Scale, 10 μm.

**Fig. S5.**
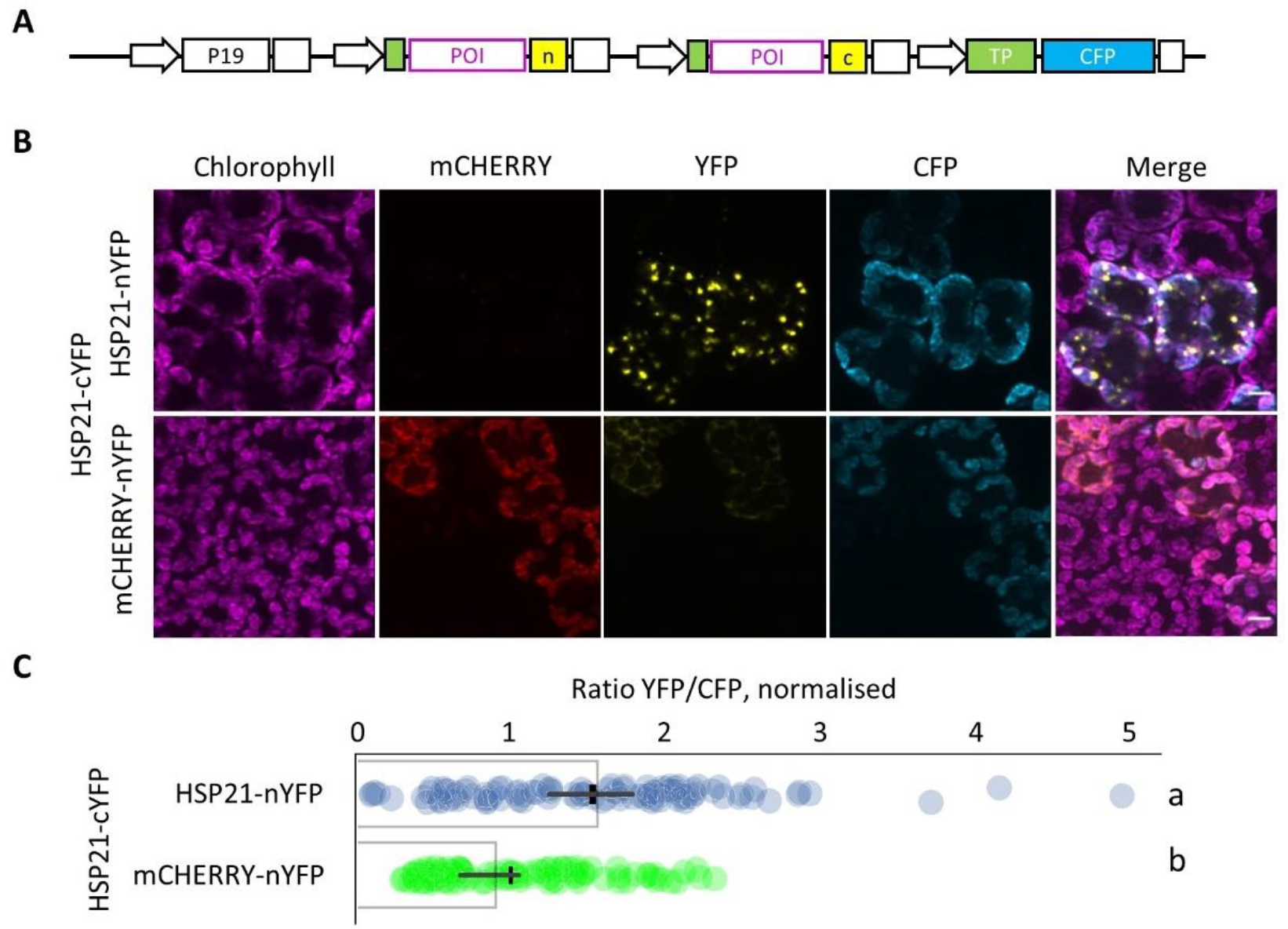
Chloroplastic CFP is unreliable as a reference FP. (a) Two different HSP21 pairs with the chloroplastic CFP reference FP were used for (b) BiFC assays in *N. benthamiana* mesophyll cells. Scale, 10 μm. Standard histogram levels are indicated below each channel. (c) Normalised BiFC signals were calculated as the ratio between total eYFP and CFP fluorescence and normalised to HSP21-cYFP / mCHERRY-nYFP. Bar indicates mean and cross indicates median +/− 95% confidence interval (n=100 transformed cells). Significance was calculated using the Kruskal Wallis test, groups are indicated by lower case letters (*P*<0.01).

**Fig. S6.**
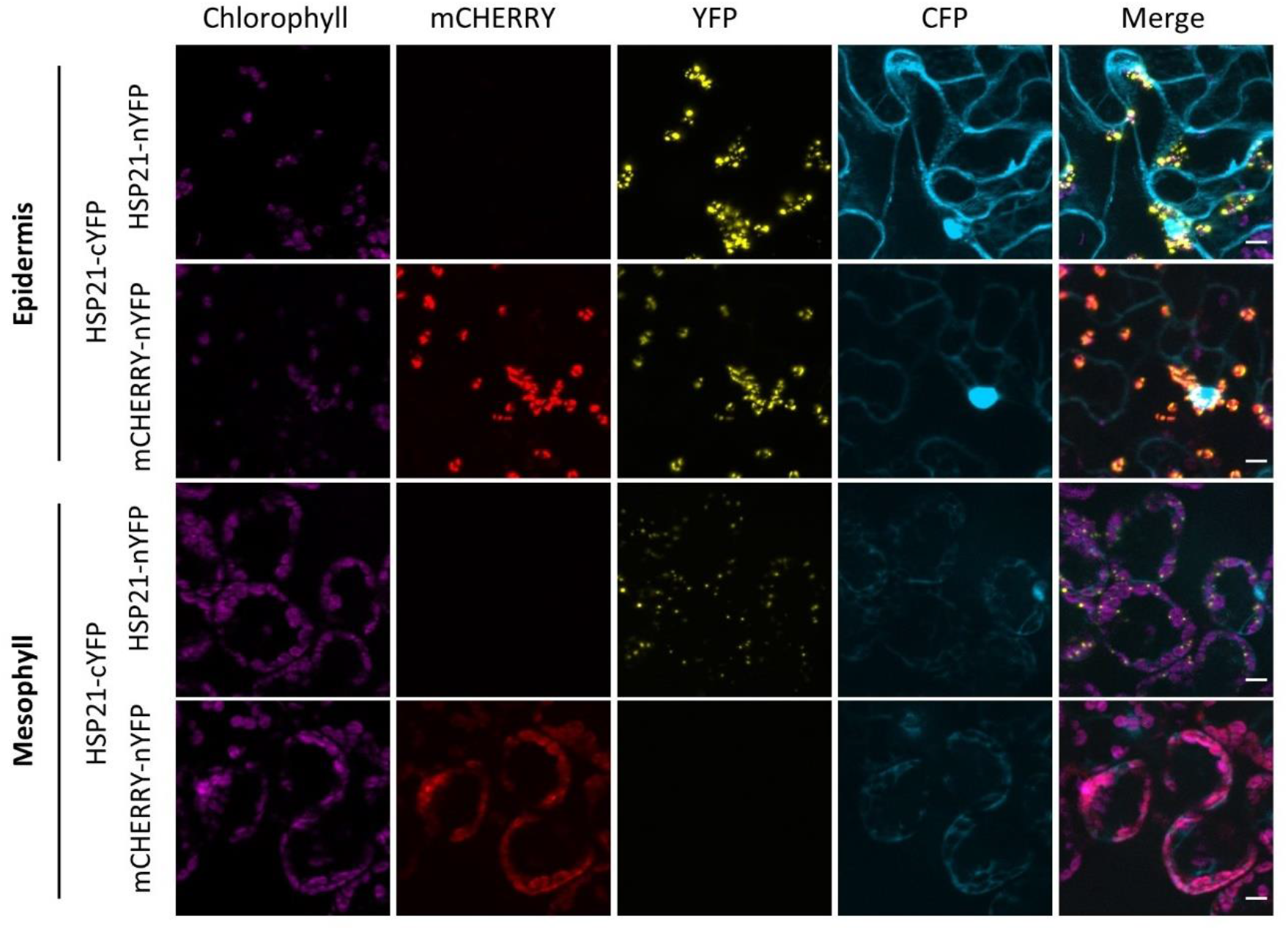
Epidermal cells have a high rate of false positive signals. HSP21-HSP21 and mCHERRY-HSP21 BiFC assays in *N. benthamiana* epidermal cells (upper two rows) and mesophyll cells (lower two rows). Mesophyll images are of the same region as shown in Fig. 2. Scale, 10 μm.

**Fig. S7.**
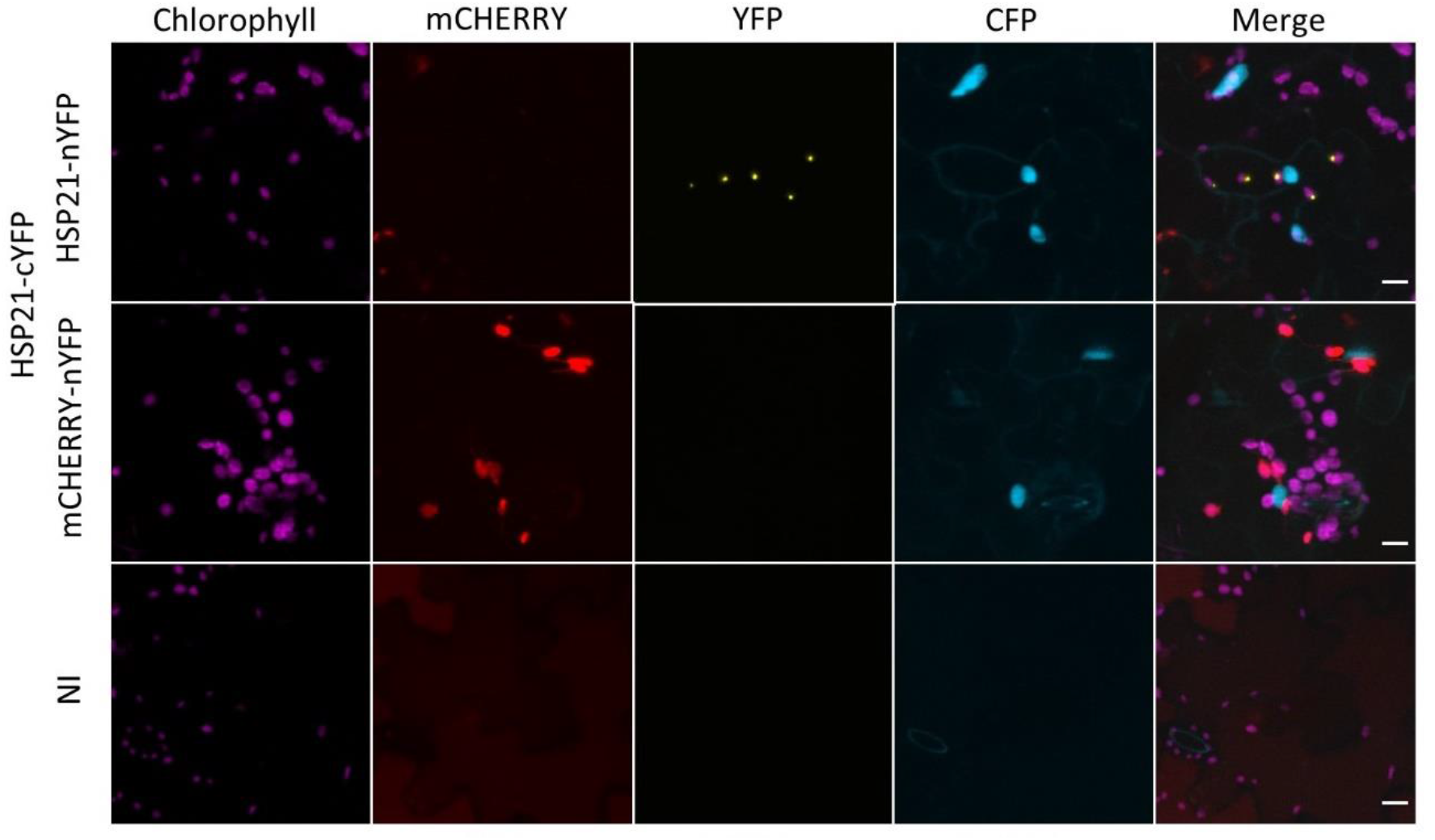
BiFC in Arabidopsis. HSP21-HSP21 and mCHERRY-HSP21 BiFC assays in *A. thaliana* epidermal cells. NI, not inoculated; scale, 10 μm.

**Fig. S8.**
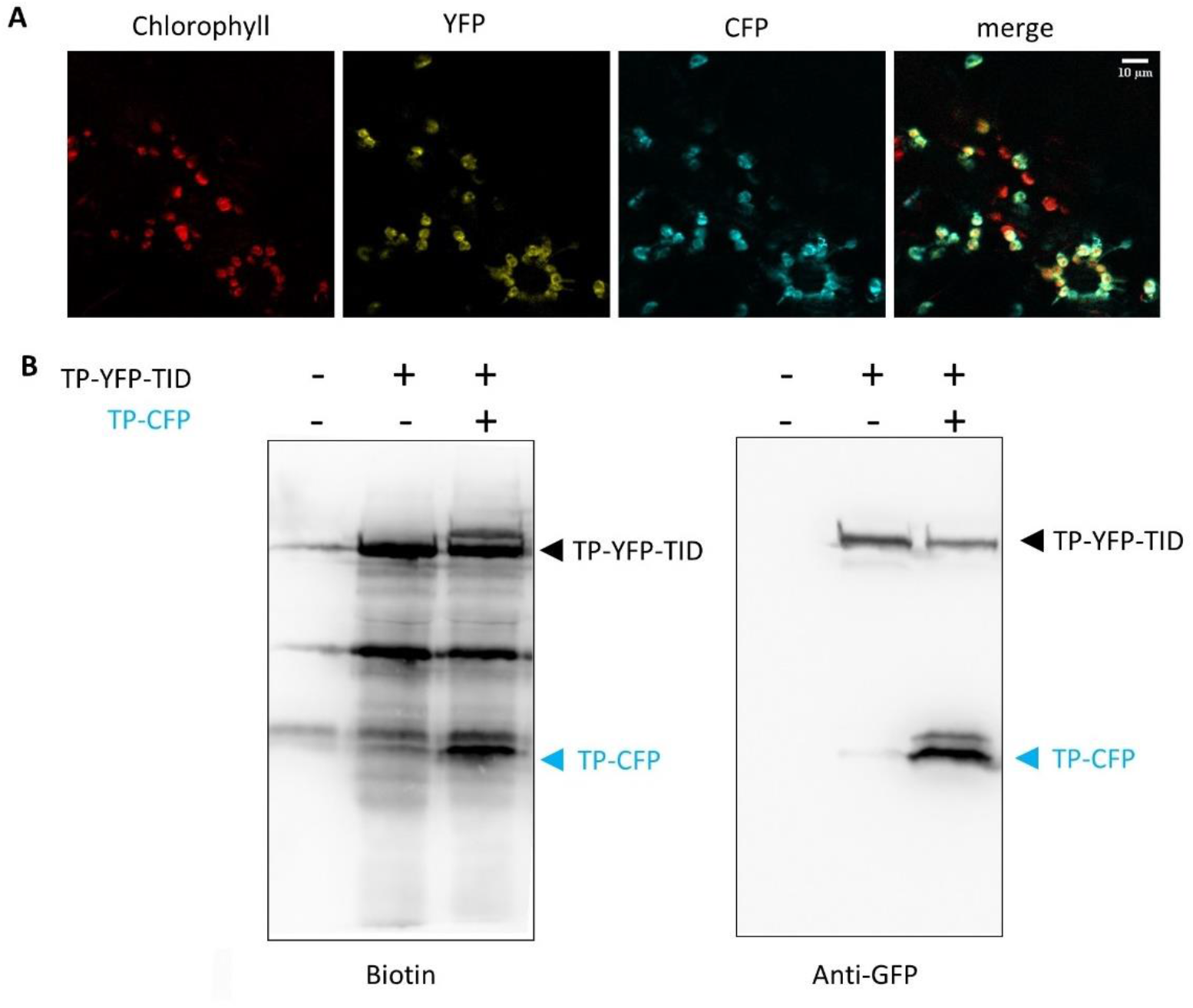
Functional test of a chloroplast TurboID module. (a) Localisation of TP-YFP-TID and TP-CFP in the chloroplasts of agro-inoculated *N. benthamiana.* Scale, 10 μm. (b) Detection of biotinylated proteins and CFP/YFP in total protein extracts. Leaf discs from non-inoculated plants, plants inoculated with TP-YFP-TID only and TP-YFP-TID + TP-CFP were incubated with biotin for 1 hr before protein extraction, separation and detection. Biotinylation of the chloroplast localised CFP (TP-CFP) was observed.

